# Artificial Intelligence for Genomics and Bioinformatics: Is Africa Lagging Behind?

**DOI:** 10.1101/2025.08.27.668885

**Authors:** Sofonias K. Tessema, Collins K. Tanui, Harris Onywera, Alan Christoffels, Nebiyu Dereje, Mosoka Papa Fallah, Yenew Kebede

## Abstract

Africa’s diverse pathogen and human genetics present significant opportunities for genomic medicine research using artificial intelligence (AI). We surveyed 250 genomics researchers across 46 of the 55 African Union Member States to assess the state of AI use. The survey revealed that only 4% of respondents reported high familiarity with the application of AI for genomics and bioinformatics.

## Genomics in Africa: Progress Amid Persistent Disparities

Africa continues to face a significant burden of recurrent and emerging infectious disease outbreaks, such as Ebola, mpox, and cholera, as well as the escalating threat of antimicrobial resistance (AMR), which causes high morbidity and mortality while straining already limited healthcare systems. Response to many of these public health events is heterogeneous, often delayed or reactive^1^. These challenges underscore the urgent need for enhanced genomic surveillance and deployment of AI to facilitate early detection, real-time tracking, forecasting, resource allocation efficiency, and informed response to pathogen threats across the continent.

The COVID-19 pandemic catalyzed the rapid expansion of genomic surveillance capacity, from 7 countries in 2019 to 41 in 2024 with functional local genomics capacity^2^, which has enabled the discovery of the clade Ib mpox variant and the application of genomics for the detection and characterization of various outbreaks in Africa^3^. However, full integration of genomic surveillance into public health systems is slow due to limited workforce, and limited access to data infrastructure, bioinformatics tools, data sharing and reporting challenges^4^. Continental efforts, such as the Africa CDC’s Pathogen Genomics Initiative (Africa PGI), are increasingly focused on bridging these gaps by fostering multi-pathogen surveillance and building technical capacity across African Union member states^5^.

Beyond pathogens, Africa’s genomic landscape reveals broader disparities in human genomics research and innovation. Despite the identification of over five million novel genetic variants in African genomes—78% of which are population-specific (Fan et al., 2023)—less than 1% of analyzed human genomes in global studies originate from the continent. This underrepresentation not only limits scientific discovery but also perpetuates algorithmic bias in AI models trained predominantly on non-African datasets.

## Artificial Intelligence: A Catalyst for Transforming Genomics and Bioinformatics in Africa

Artificial intelligence holds transformative potential for genomics and bioinformatics in Africa, offering tools for rapid data processing, analyses and interpretation, predictive analytics, and enhanced epidemiological insights. AI-driven platforms are increasingly recognized for their ability to streamline pathogen detection, track mutations in near real-time, and support early outbreak response—functions critical in resource-limited settings^6^. The COVID-19 pandemic demonstrated that, when equipped with sufficient tools and resources, African scientists can achieve scientific breakthroughs with global relevance^7^. Capacity-building efforts, like those spearheaded by the Africa PGI, are training a new generation of African bioinformaticians and genomic epidemiologists to harness genomics tools for public health^8,9^. There is anecdotal to limited published data on AI knowledge and application in genomics and bioinformatics in Africa. Here, we conducted a continental survey to capture the current status, knowledge, adoption, challenges, and limitations of AI applications for genomics and bioinformatics in Africa.

## Methods

### Study Design, Data Collection, and Preprocessing

We conducted a cross-sectional, continent-wide survey to assess the awareness, adoption, and perceived barriers to artificial intelligence (AI) applications in genomics and bioinformatics across Africa. The survey was developed by a multidisciplinary team at Africa CDC’s Centre for Laboratory Diagnostics and System through the Africa Pathogen Genomic Initiative (Africa PGI) and disseminated electronically between March and May 2025 through institutional mailing lists, regional networks, and professional platforms targeting genomics and public health professionals.

The structured questionnaire included both closed- and open-ended items designed to capture demographic characteristics, institutional affiliations, AI knowledge levels, current use cases, perceived utility, and training needs. The instrument was piloted with a small group of genomics experts to ensure clarity, contextual relevance, and technical accuracy. Participation was voluntary, and only responses from individuals who provided informed consent were included in the final analysis.

A total of 250 respondents from 46 African Union Member States completed the survey. Descriptive statistics were used to summarize the data. Likert-scale responses were visualized using bar charts and Sankey diagrams to illustrate trends in AI familiarity and application. All data were anonymized prior to analysis to ensure confidentiality

## Results and Discussions

A continent-wide survey of 250 genomics professionals from 46 African countries reveals a significant gap in AI awareness, preparedness and adoption for genomics and bioinformatics. Among the respondents, 59 (23.60%) hold a PhD degree; 104 (41.60%) a Master’s Degree, 72 (28.80%) a Bachelor’s Degree, and 15 individuals (6.00%) have other qualifications. A total of 136 respondents (54.40%) are affiliated with public health institutions; 92 (36.80%) are affiliated with academic and research institutions; and 22 (8.80%) are affiliated with non-governmental, private, and other institutions.

While 90% (225/250) recognized AI’s benefits and 85% viewed it as a catalytic force for genomics and bioiformatics in Africa, a critical barrier looms; only 17% reported substantial knowledge of general AI applications and a mere 4% reported high familiarity with its application in genomics and bioinformatics (Figure 1). This disparity highlights a critical capacity gap that hinders progress. Addressing this issue through targeted and accessible trainings and mentorship is vital to fully leverage AI’s potential in advancing public health genomics across Africa^2,9,10^.

**Figure 1.**
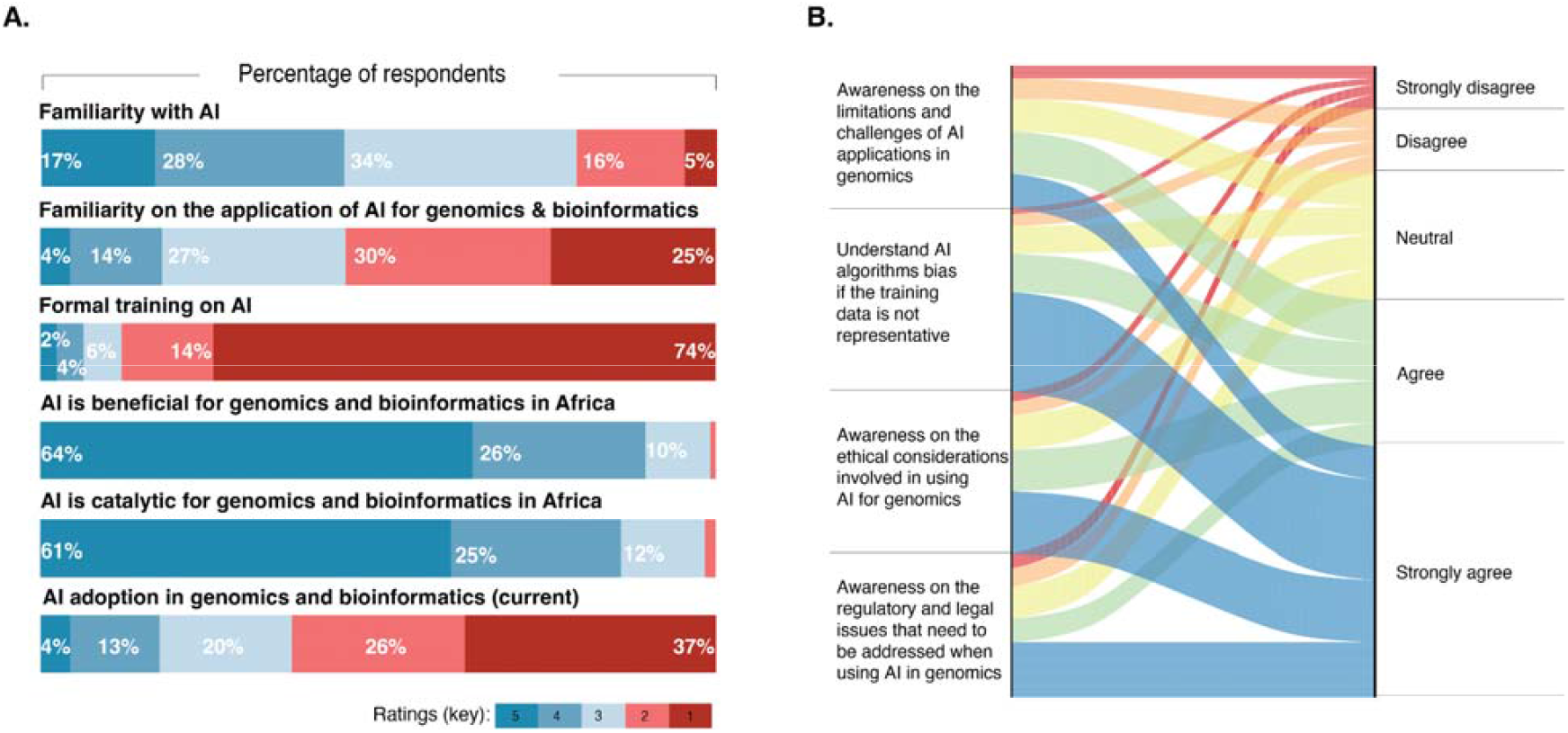
Knowledge, preparedness, adoption and limitations of AI applications in genomics and bioinformatics. **(A)** Bar chart illustrating the distribution of responses (n = 250) across six key dimensions. Responses are color-coded on a Likert scale from 1 (dark red) to 5 (dark blue). **(B)** Sankey diagram depicting the flow of respondents’ awareness levels across four critical considerations in AI use for genomics.

Building on this, the survey further reveals a strong and widespread demand for training in AI applications relevant to genomics and bioinformatics. An overwhelming 90.4% (226/250) of respondents expressed interest in acquiring AI-relevant skills. The most sought-after areas include genomic data processing, analysis, and interpretation (47%), followed by predictive modeling (18.7%), machine learning (17%), and ethical considerations (16%). However, despite this enthusiasm, only 0.8% (2/250) have enrolled and completed a formal AI training, and just 6% (16/250) have received AI related training in the past—leaving 93.8% of respondents without foundational expertise.

The barriers to training are substantial and disproportionately affect African countries. The lack of available training programs (78%) and the high cost of training (60%) are the most frequently cited obstacles, exacerbating existing inequities in access to advanced genomic and bioinformatics tools. These findings paint a picture of a continent poised for a genomic revolution yet constrained by a critical skills deficit. The emphasis on data analysis reflects the urgent need to manage and interpret large-scale genomic datasets, while the demand for ethical and regulatory related trainings underscores a growing awareness of the importance of safeguarding Africa’s genomic sovereignty. This collective aspiration emphasizes the need for locally tailored, equity-focused AI training initiatives that ensure Africa can responsibly and effectively harness its rich genetic diversity on the global stage.

### Slow adoption of AI in Africa

Despite strong interest, AI adoption in Africa remains limited. Only 4% of surveyed institutions have fully integrated AI tools such as machine learning algorithms and natural language processing, with most implementations concentrated in laboratories equipped with robust sequencing capacity. The current use cases—primarily in bioinformatics pipelines (60%) and variant identification (50%)—are consistent with global trends and demonstrate a focused, albeit narrow, application of AI in public health.

Encouragingly, 85% of respondents indicated plans to adopt AI in the near future—a clear signal of intent. Yet significant barriers threaten to stall progress. The major among the deficits are training (80%) and infrastructure (70%), which limit broader deployment. Financial limitations (55%) and inconsistent regulatory frameworks (60%) also highlight the urgent need for African-led governance structures that can guide the responsible and context-specific implementation of AI for genomics and bioinformatics.

### AI is viewed as a game-changer

Our survey reveals strong confidence in the transformative potential of AI in public health genomics and bioinformatics across Africa. Respondents rated AI’s potential and utility highest in data analysis (84%), early disease detection (82%), and diagnostic accuracy (80%). These findings highlight AI’s perceived value in enhancing pathogen surveillance and enabling rapid outbreak response—two areas that are critical for strengthening public health systems on the continent and the broader global health security.

In contrast, lower ratings for applications such as report writing (60%) and resource allocation (58%) suggest that AI is not yet widely seen as a tool for supporting administrative or operational functions. This distinction reflects the continent’s immediate priorities: rapid variant identification and outbreak forecasting, which demand advanced analytical capabilities over back-office automation. However, enthusiasm for AI is tempered by legitimate concerns. Sixty-five percent of respondents cited worries about algorithmic bias—pointing to a broader challenge: the underrepresentation of African genomic data in global AI training datasets. Addressing this imbalance is essential to ensure that AI systems are not only effective but also equitable in serving Africa’s diverse populations.

### Bridging the Gaps: the Leapfrogging Potential of AI in African Genomics

Despite recent progress, African genomics faces persistent challenges that hinder its scalability and integration into public health systems. Acute shortage of genomics and bioinformatics experts, which limits the ability of institutions to conduct robust, real-time analyses and public health insights^9,11^. Compounding this issue are critical data data infrastructure and analytics gaps, often depending on external partners for data analysis and interpretation, thus delaying outbreak response. Furthermore, fragmented and inefficient data-sharing mechanisms continue to obstruct cross-border collaboration and limit the continent’s collective genomic intelligence^4^. Nonetheless, AI presents a unique opportunity to leapfrog these systemic challenges by enabling broader participation in genomics through intuitive, user-friendly, and AI-powered analytical tools. Such technological democratization, coupled with strengthened regional networks and data governance frameworks, could transform Africa’s capacity to lead in the global genomics arena.

### Conclusion and way forward: A call for an AI-empowered Genomics in Africa

Africa is on the brink of a genomic revolution, yet a critical gap in genomics infrastructure, workforce, and AI capacity threatens to leave the continent behind. Despite overwhelming interest and recognition of AI’s transformative potential in genomics and bioinformatics, African public health and genomics researchers face steep barriers—limited training, inadequate infrastructure, and underrepresentation in global datasets.

We call on governments, philanthropists, public health agencies, and local communities to act now. Governments must invest in national data and AI strategies and governance that prioritize genomic research and integrate AI into public health systems. Philanthropic organizations should continue to catalyze inclusive, affordable training programs to build a skilled workforce. National Public Health Institutions must champion ethical, African-led AI governance frameworks to ensure equitable and context-specific implementation.

The public health and scientific imperatives are clear: AI can help Africa leapfrog traditional barriers, enabling real-time outbreak detection, personalized medicine, stronger health systems, and innovations that not only serve Africa but the global health security. Urgent, coordinated action is needed to include African in this transformative wave. Let us unite to build a future where Africa leads in genomic innovation—powered by AI, driven by equity, and rooted in local leadership. The time to act is now.

## References

1. Bosa, H. K. et al. How to prepare for the next inevitable Ebola outbreak: lessons from West Africa. Nat Med 30, 3413–3416 (2024).

2. Mboowa, G. et al. Africa in the era of pathogen genomics: Unlocking data barriers. Cell 187, 5146–5150 (2024).

3. Vakaniaki, E. H. et al. Sustained human outbreak of a new MPXV clade I lineage in eastern Democratic Republic of the Congo. Nat Med 30, 2791–2795 (2024).

4. Inzaule, S. C., Tessema, S. K., Kebede, Y., Ogwell Ouma, A. E. & Nkengasong, J. N. Genomic-informed pathogen surveillance in Africa: opportunities and challenges. Lancet Infect Dis 21, e281–e289 (2021).

5. Mboowa, G. et al. The rise of pathogen genomics in Africa. F1000Res 13, 468 (2024).

6. Tanui, C. K., Ndembi, N., Kebede, Y. & Tessema, S. K. Artificial intelligence to transform public health in Africa. Lancet Infect Dis 24, e542 (2024).

7. Tegally, H. et al. The evolving SARS-CoV-2 epidemic in Africa: Insights from rapidly expanding genomic surveillance. Science 378, eabq5358 (2022).

8. Onywera. How to sustain a public-health genomics and bioinformatics workforce in Africa.

9. Onywera, H. et al. Boosting pathogen genomics and bioinformatics workforce in Africa, has been accepted for publication. The Lancet Infectious Diseases e281– e289 (2023).

10. Pronyk, P. M. et al. Advancing pathogen genomics in resource-limited settings. Cell Genom 3, 100443 (2023).

11. Aruhomukama, D., Galiwango, R., Meehan, C. J. & Asiimwe, B. Enhancing genomics and bioinformatics access in Africa: an imperative leap. The Lancet Microbe 5, e410–e411 (2024).

